# Astrocytes regulate neuronal network burst frequency through NMDA receptors species- and donor-specifically

**DOI:** 10.1101/2023.12.09.570906

**Authors:** Noora Räsänen, Jari Tiihonen, Marja Koskuvi, Šárka Lehtonen, Nelli Jalkanen, Nelli Karmila, Isabelle Weert, Olli Vaurio, Ilkka Ojansuu, Markku Lähteenvuo, Olli Pietiläinen, Jari Koistinaho

**Affiliations:** Neuroscience Center, University of Helsinki; Helsinki, Finland; Department of Clinical Neuroscience, Karolinska Institutet, and Center for Psychiatric Research, Stockholm City Council, Stockholm, Sweden; A.I. Virtanen Institute for Molecular Sciences, University of Eastern Finland, Kuopio, Finland; Department of Forensic Psychiatry, University of Eastern Finland, Niuvanniemi Hospital, Kuopio, Finland; Helsinki Institute of Life Science, University of Helsinki; Helsinki, 00014; Finland; Drug Research Program, Division of Pharmacology and Pharmacotherapy, University of Helsinki, FI 00014 Helsinki, Finland

## Abstract

**Background:** Development of synaptic activity is a neuronal key characteristic that relies largely on interactions between neurons and astrocytes. Although astrocytes have known roles in regulating synaptic function and malfunction, the use of human or donor-specific astrocytes in disease models is still rare. Rodent astrocytes are routinely used to enhance neuronal activity in cell cultures, but less is known how human astrocytes influence neuronal activity.

**Methods:** We established human induced pluripotent stem cell (hiPSC)-derived neuron-astrocyte co-cultures and studied their functional development on microelectrode array (MEA). We used cell lines from 5 neurotypical control individuals and 3 pairs of monozygotic twins discordant for schizophrenia. A method combining Ngn2 overexpression and dual SMAD inhibition was used for neuronal differentiation. The neurons were co-cultured with hiPSC-derived astrocytes differentiated from 6-month-old astrospheres or rat astrocytes.

**Results:** We found that the hiPSC-derived co-cultures develop complex network bursting activity similarly to neuronal co-cultures with rat astrocytes. However, the effect of NMDA receptors on neuronal network burst frequency (NBF) differed between co-cultures containing human or rat astrocytes. By using co-cultures derived from patients with schizophrenia and unaffected individuals, we found lowered NBF in the affected cells. We continued to demonstrate how astrocytes from an unaffected individual rescue the lowered NBF in the affected neurons by increasing NMDA receptor activity.

**Conclusions:** Our results indicate that astrocytes participate in the regulation of neuronal NBF through a mechanism involving NMDA receptors. These findings shed light on the importance of using human and donor-specific astrocytes in disease modeling.

## Introduction

Human induced pluripotent stem cell (hiPSC)-derived brain cells provide an accessible model to study mechanisms of brain disorders in a human context. Neuronal differentiation protocols based on NGN2 transcription factor overexpression have been widely adopted to rapidly generate homogenous populations of excitatory neurons (1,2). These fast protocols have provided an advantage over traditional differentiation methods in terms of scalability and homogeneity of neuronal cultures. In addition, new protocols coupling the NGN2 expression with small molecule patterning have further improved the subtype specificity of the neurons (2,3). These features have improved the applicability of hiPSC-derived neurons for disease modeling.

Neuronal synaptic maturation depends on surrounding astrocytes that actively regulate neuronal properties (2,4). To enhance the synaptic and functional maturation of the NGN2-expressing neurons *in vitro*, the neurons are often co-cultured with rodent astrocytes. This mixed-species model promotes rapid development of neuronal network-level activity already in 4 weeks of differentiation (2,5). Despite the effectiveness of rodent astrocytes in supporting neuronal maturation, they differ from human astrocytes in terms of structural complexity, subtype diversity, developmental timeline, metabolic flexibility, response to inflammatory conditions and many other features (6–8). This has raised a question to what extent rodent astrocytes can substitute human glial cells and recapitulate features that are relevant for human neural functions. Furthermore, an increasing number of studies has shown that astrocytes have a role in the pathogenesis of developmental (4,9) and neurodegenerative disorders (10,11). This has evidenced the need for fully human neuron-astrocyte co-cultures for modeling development and disorders of the human brain. However, little is known how hiPSC-derived astrocytes influence neuronal activity and whether the variance in these properties in neurons is influenced by the astrocyte donor.

The aim of this study was to develop a hiPSC-derived neuron-astrocyte co-culture model to recapitulate both neuron and astrocyte-related functional alterations in brain diseases. First, we characterized the functional development of hiPSC-derived neuron-astrocyte co-cultures across time. We compared the fully human co-cultures to mixed-species co-cultures of human neurons and rat astrocytes. Finally, using cell lines from individuals with treatment-resistant schizophrenia (TRS), we prepared co-cultures for studying the role of the different cell types in regulating neuronal network-level activity in schizophrenia.

## Methods and materials

### Cell lines

hiPSC lines were derived from 5 neurotypical control individuals and 3 pairs of monozygotic twins discordant for TRS. The patients with schizophrenia had a history of clozapine use (Table S1). The reprogramming and characterization of the hiPSC lines have been previously documented (12). All participants provided written informed consent, and the work has been approved by the Ethics Committee of the Helsinki University Hospital District, license number 262/EO/06.

### Neuronal differentiation

Neuronal differentiation was carried out by Ngn2 overexpression coupled with dual SMAD and WNT inhibition, as previously described (2), to generate homogenous populations of cortically patterned glutamatergic neurons (Figure 1A). The hiPSCs were infected overnight with 3 lentiviruses: Tet-O-NGN2-PURO, Tet-O-FUW-eGFP and FudeltaGW-rtTA (>10^9^ IFU/ml, Alstem), using MOI = 10. The differentiation was initiated by adding 2 μg/ml Doxycycline hyclate (2431450, Biogems) in Essential 8 medium (A15169-01, Gibco). Doxycycline was used in the cell culture medium from this day onward. On day 1, N2 medium (DMEM/F-12 (21331-020, Gibco), 1:100 Glutamax (35050-038, Gibco), 1:100 N2 supplement (15410294, Gibco, 1:67 20% Glucose) was prepared and supplemented with 10 μM SB431542 (SB, S4317, Sigma), 100 nM LDN193189 (LDN, SML0559, Sigma), 2 μM XAV939 (2848932, Biogems) and 2 mg/ml Doxycycline. On day 2, the medium was changed to N2 medium supplemented with 1:2 of the day 1 supplements as well as 5 μg/ml puromycin (100552, MP biomedicals) to select for infected cells. On day 3, the cells were given the same media as on day 1. The final plating of the neurons was done on day 4. The cells were detached with Accutase (11599686, Gibco) for 5 min and centrifuged at 300 rcf for 4 min before counting. The cells were plated on wells coated with 50 μg/ml Poly-L-ornithine (Sigma, P3655) and 10 μg/ml laminin (Sigma, L2020). Neurons were plated 1:1 with astrocytes using 60 000 neurons/cm^2^ for immunocytochemistry (ICC) and 60 000 neurons/well for microelectrode array (MEA; Supplementary Figure 1). Neurons were fed with Neurobasal medium (NBM, (21103-049, Gibco), 1:100 Glutamax, 1:200 MEM NEAA (11140-035, Gibco), 1:67 20% Glucose, 1:50 B27 without vitamin A (12587001, Gibco) supplemented with 10 ng/ml BDNF (450-02, PeproTech), 10 ng/ml GDNF (450-10, PeproTech) and 10 ng/ml CNTF (450-13, PeproTech) from day 4 onward. On days 7-8, proliferating cells were eliminated from the cultures with 10 μM FUDR (4659, Tocris). Half of the medium was changed three times a week.

**Figure 1:**
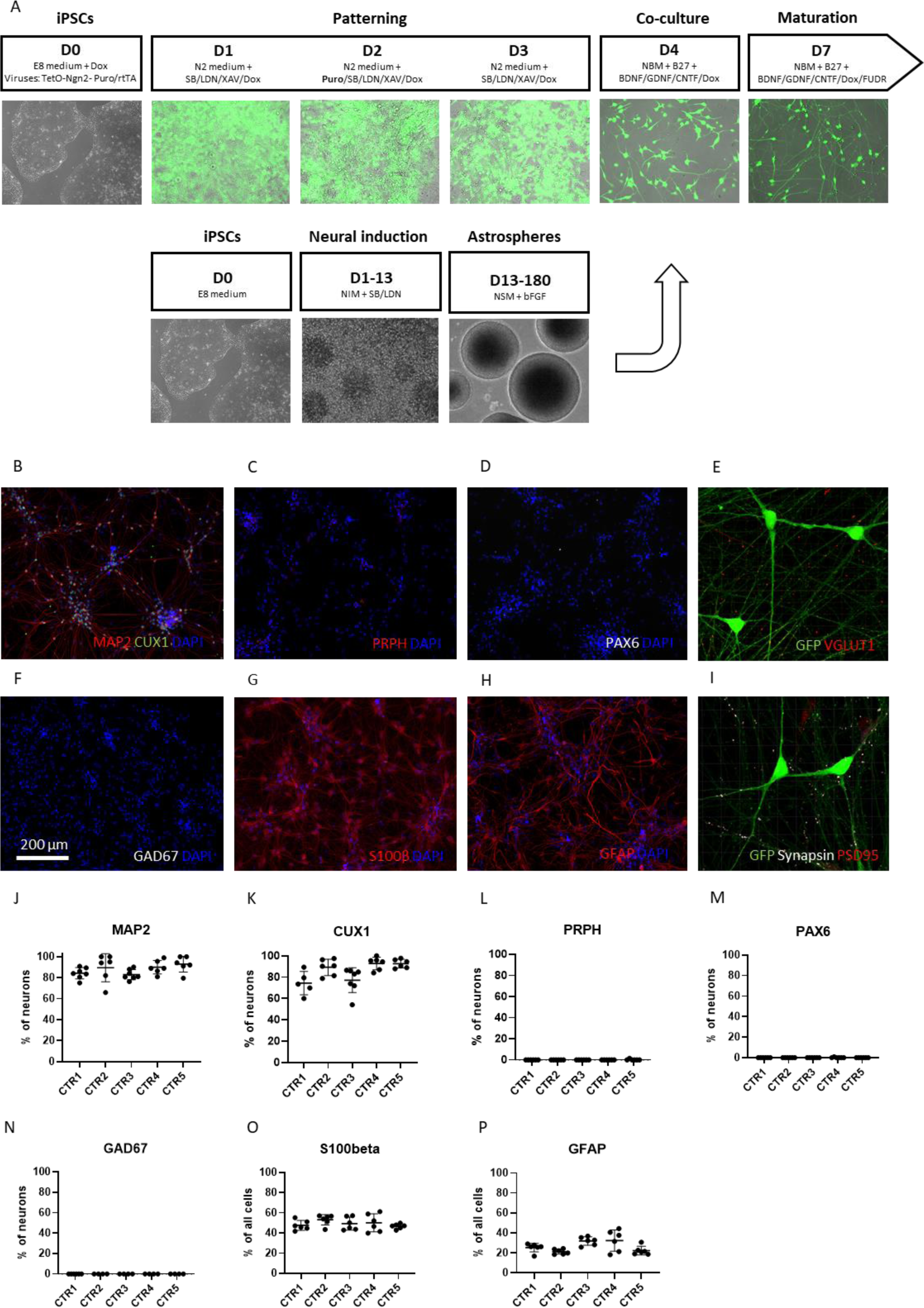
Characterization of hiPSC-derived neuron-astrocyte co-cultures. A. Differentiation of neurons and astrocytes. B, J-K. 5-week-old neurons expressed neuronal marker MAP2 and superficial cortical layer marker CUX1. C-D, L-M. The cultures did not stain for PRPH and PAX6, indicating the absence of peripheral neurons and differentiation-resistant PAX6-positive NPCs. E. The neurons expressed excitatory synaptic marker VGLUT1. F, N. There were no GAD67-positive cells in the cultures indicating the absence of GABAergic neurons. G-H, O-P. Approximately 50% of the cells in the co-cultures were S100β-positive astrocytes and 8-20-30% of the cells expressed astroglial marker GFAP. I. Neurons expressed co-localized Synapsin and PSD95 proteins for juxtaposed pre- and postsynaptic puncta, respectively. This indicated the presence of structural synapses. (n = 5 cell lines, data was collected from 3 independent experiments)

### Astrocyte differentiation

The astrocyte differentiation was done as previously described (9,13). Briefly, neural induction was performed by culturing hiPSCs with 10 μM SB and 2 μM LDN. After 10 days of induction, NPCs arranged in rosettes were manually picked and transferred to ultra-low attachment plates, where the NPCs formed spheres. These spheres were cultured and expanded with bFGF (100-18B, Peprotech) for 6-9 months, after which they were dissociated and plated in co-cultures with neurons at 1:1 density. Astrocytes from different cell lines were cultured concurrently and represented the same age when plated. The astrocytes used in this study have been fully characterized before (9).

### Rat primary astrocytes

Rat cortices were extracted from Wistar rat embryos at E17-18 as previously described (14). After initial cell preparation, the purified cortical cells were plated on T75 flasks with 20 ml DMEM High Glucose (ECB7501L, Euro Clone) supplemented with 10 % fetal bovine serum (10500-064, Gibco), 1 % L-Glutamine (BE17-605E, BioWhittaker) and 1 % Penicillin/Streptomycin (DE17-602E, BioWhittaker). After 7-8 days of culturing, the astrocytes reached confluency, and contaminating microglia and oligodendrocytes were removed by shaking the flask at 240 rpm for 6h as previously described (15). The cells were split using Trypsin (0.05%)-EDTA and the media was changed once every two weeks. The rat astrocytes were plated with hiPSC-derived neurons 1:1 using 60 000 astrocytes/well for MEA.

### Immunocytochemistry

For ICC staining, 5-week-old neurons were fixed with 4% formaldehyde for 20 min and washed twice with PBS. The cells were permeabilized with 0.25% Triton X-100 (T8787, Merck) in PBS for 1 hour. Unspecific binding sites were blocked using 5% normal goat serum (NGS, S26, Merck) for 1 hour. Primary antibodies against MAP2 chicken (1:500, Abcam, Ab92434), CUX1 mouse (1:500, Abcam, ab54583), VGLUT1 rabbit (1:300, Sigma, vo389-200), GAD67 mouse (1:500, Abcam, ab26116), PRPH rabbit (1:2000, Novus, NB300-137), PAX6 rabbit (1:500, Thermo, 42-6600), Synapsin Oyster 650 mouse (1:500, Synaptic Systems, 106011C5), PSD95 rabbit (1:500, Cell Signaling Technologies, 3450), GRIN1 rabbit (1:500, Cell Signaling Technologies, 5704), s100β rabbit (1:100,Abcam, 52642) and GFAP rabbit (1:500, DAKO, Z033429-2) were used. The primary antibody mixture was prepared in 5% NGS and incubated overnight at 4°C on a shaker. The cells were washed 3 x with PBS. Secondary antibodies including Goat anti-chicken Alexa Fluor 568 (A11041), Goat anti-rabbit Alexa Fluor 568 (A11011), Goat anti-mouse Alexa Fluor 633 (1:400, A21052, all from Thermo Fisher) were used. The secondary antibody mixture was prepared in PBS and incubated for 2 hours with the cells. Finally, the cells were washed 2 x with PBS and 1 x with DAPI (1:2000, Sigma) for 10 min. The samples were mounted using Fluoromount-G (00-4958-02, Thermo Fisher) and imaged using EVOS M5000 fluorescence microscope or Andor Dragonfly spinning disk confocal microscope (Nikon).

### Image analysis

The image analysis for neuronal characterization was done with ImageJ (NIH). The neurons were separated from the astrocytes during the analysis based on their GFP expression, and the astrocytes were identified based on S100β expression. The images were thresholded using the default option to select for the cells expressing the target protein. The image calculator was used to co-localize the selected cells with a reference marker. DAPI was used as a reference for all cells and GFP was used as a reference for neurons. The analyze particles feature was used to quantify the number of cells expressing the proteins of interest. Analysis of the 3D images was performed with Imaris 10 (Bitplane). The Surfaces function was used to select the cells for analysis in order to measure the mean intensity of the staining inside the surface.

### MEA recordings

The electrophysiological activity was recorded with Maestro Edge MEA system using AxIS Navigator software and 24-well CytoView plates containing 16 electrodes (Axion Biosystems). The recordings were performed at 37 °C temperature in a 5% CO_2_ atmosphere. To start the measurements, the well plate was placed in the MEA system and the temperature and CO_2_ were allowed to stabilize for 10 min. The baseline activity was measured for 10 min. For pharmacological tests, 10 μM NBQX (N138, Sigma), 25 μM D-AP5 (A8054, Sigma), 100 μM GABA (Sigma), 10 μM Ifenprodil (I2892, Sigma) and 3 μM TCN-201 (SML0416, Sigma) were used according to previous studies (2,12,16,17). The baseline activity was recorded for 10 min prior to treatment. After this, the pharmacological treatments were carried out by pipetting 5 μl of each compound into wells containing 500 μl of culture media. The plate was then placed back in the MEA device and incubated for 10 min before a 10 min recording was started.

### MEA data analysis

The AxIS Navigator software was used for spike sorting during the recordings. The spike threshold was set to 5 x the standard deviation of the estimated noise. The burst detection was done using Neural Metric Tool (Axion biosystems). The minimum number of spikes per burst was set to 5 and the maximum inter-spike interval (ISI) within a burst was set to 100 ms. For NB detection, the Envelope algorithm was used due to its ability to merge repetitive sub-bursts within a network event into a single burst. A threshold factor value of 2 and a minimum inter-burst interval (IBI) 100 ms were selected for the analysis. The minimum number of electrodes in a NB was set to 25 % and a burst inclusion value was set to 75%. The Envelope algorithm was not suitable for the analysis of NBQX-treated samples that contained a great amount of non-synchronous bursting activity. Instead, we used Adapted algorithm that more efficiently separated the NBs from non-synchronous bursts. The minimum number of spikes per NB was set to 50 and the minimum number of electrodes in NB was set to 25 %. The NBs in the patient lines were analyzed using the max ISI algorithm that performed the best in terms of defining NB duration in samples with short bursts and bursting outside the NBs. The max ISI value was set to 50 ms and the minimum number of spikes per NB was set to 50.

NeuroExplorer (Plexon) software was used to analyze high frequency bursting activity in the samples. First, spike-sorted .spk files generated by the AxIS Navigator software were converted to rate histograms using a 0.025 s bin width in NeuroExplorer. Electrodes with fewer than 5 spikes per minute were removed from the files. Power spectral densities (PSD) were drawn for each electrode using Welch’s windowing function. The same function was used for drawing the spectrograms. For visualization of spectrograms and PSDs, Log of PSD (dB) normalization was used, and for the analysis of high frequency bursting, raw PSD values were used. The power of the signal at frequencies 0.5-4 Hz, 4-8 Hz and 8-12 Hz was acquired from the PSDs for each electrode by averaging values within the defined frequency range. Finally, the values for each sample were acquired by averaging the values from the electrodes. The results for the power of high frequency bursting were presented as a ratio to the network burst frequency (NBF) in each well.

### qRT-PCR

RNA was extracted from neuron-astrocyte co-cultures at 5 weeks using RNeasy Mini kit (74104, Qiagen) following the manufacturer’s instructions. The extracted RNA was eluted in nuclease-free water. The cDNA conversion was performed with Maxima reverse transcriptase enzyme approach using Random hexamer primer (S0142, Fermentas), 10 mM dNTP (R0192, Fermentas), 40 U/μl Ribonuclease inhibitor (E00381, Fermentas) and Maxima reverse transcriptase (EPO742, Fermentas). For the qPCR reaction, Maxima Probe qPCR Master Mix (K0261, Thermo Fisher Scientific) and primers for GRIN1 (Hs00287446_m1 and Rn01436030_m1, Thermo Fisher Scientific), GRIN2A (Hs00168219_m1, Thermo Fisher Scientific) and GRN2B (Hs01002013_m1, Thermo Fisher Scientific) were used. The gene expression was normalized to GAPDH (Hs99999905_m1, Thermo Fisher Scientific) or ACTB (Hs99999903_m1 and Rn00667869_m1, Thermo Fisher Scientific) using Q-gene program (18).

### Statistical analysis

The statistical analysis was performed with GraphPad Prism 9.4.1 and RStudio 2022.12.0. Mann-Whitney U test was used to compare the differences between time points or co-cultures containing astrocytes from different species or donors. Paired t test was used for the investigation of pharmacological responses in neurons using GraphPad Prism. The normal distribution of the data was verified with Kolmogorov-Smirnov test. p-values for the comparison of cells from affected and unaffected individuals were derived from ANOVA using general mixed linear regression model in the lme package in R. The statistical tests for patient comparisons were corrected for multiple testing (21 tests) using Benjamin-Hochberg procedure implemented in GraphPad Prism.

## Results

### Establishment of human iPSC-derived neuron-astrocyte co-cultures

In order to study neuronal maturation in human neuron-astrocyte co-cultures, we generated cultures of hiPSC-derived excitatory neurons and astrocytes from 5 neurotypical control individuals using previously published protocols (2,13). The neurons and astrocytes were plated together after 4 days of neural induction (Figure 1A) and the maturation of the co-cultures was continued until 5 weeks of neuronal differentiation. To evaluate the maturational stage and subtype specificity of the cells, we characterized the cultures using ICC at 5 weeks of differentiation. The neurons were labeled with GFP to separate them from astrocytes during the analysis (Figure S1A). After 5 weeks, 83-93 % of the neurons expressed mature neuronal marker MAP2 (Figure 1B, J), and 71-95 % of the neurons were positive for superficial layer cortical marker CUX1 (Figure 1B, K). Hence, a great majority of the neurons had reached a mature neuronal identity and possessed features of superficial layer cortical neurons. Reassuringly, we observed minimal to no staining for contaminating cell types including peripheral neurons (PRPH), differentiation-resistant NPCs (PAX6) and GABAergic neurons (GAD67, Figure 1C-F, L-N). Instead, the neurons expressed glutamate transporter VGLUT1, indicating the presence of glutamatergic neurons (Figure 1E). In addition, we detected juxtaposed pre-and postsynaptic proteins along the neurites, which confirmed the presence of structural synapses in the co-cultures (Figure 1I).

Finally, we used staining for canonical astrocyte markers to confirm the presence of the astrocytes in the co-cultures. The astrocytes used in this study have been fully characterized before (9). The expression of astrocyte-specific markers including S100β, GFAP, ALDH1L1, Vimentin and AQP4 has been confirmed with qPCR and ICC staining. In addition, we have performed full transcriptomic profiling of the cells and confirmed their functionality with glutamate uptake assay and calcium imaging (9). After 5 weeks of maturation in co-cultures, we detected S100β expression in 47-53% of the cells, whereas 21-32 % of all cells expressed astroglial marker GFAP (Figure 1 G-H, O-P). Given that we have previously shown that approximately 100% of these astrocytes express S100β, we concluded that the percentage of astrocytes in the co-cultures remained close to the original 50%. In summary, we were able to produce homogenous co-cultures of NGN2-induced neurons and hiPSC-derived astrocytes with structural synapses. The neurons in these cultures possessed features of superficial layer cortical neurons and the neuron-astrocyte ratio remained close to the original plating ratio.

### NGN2-expressing neurons develop synchronous bursting activity in co-cultures with hiPSC-derived astrocytes

After confirming robust neuronal maturation in co-cultures with hiPSC-derived astrocytes, we set out to explore electropshysiological activity in these cultures. The functional development of NGN2-expressing neurons has been previously characterized in mono-cultures and in co-cultures with rodent astrocytes (2,5). Typically, neurons in the mixed-species co-cultures produce synchronous activity by 4 weeks of differentiation whereas the neurons in mono-cultures do not develop synchronous activity (2,5). To investigate whether NGN2-expressing neurons develop synchronous activity in co-cultures with hiPSC-derived astrocytes, we prepared donor-specific co-cultures of NGN2-expressing neurons and hiPSC-derived astrocytes from 5 cell lines for electrophysiological recordings. We compared the development of neuronal activity in the hiPSC-based co-cultures to neuronal monocultures and mixed-species co-cultures across 9 weeks of maturation on MEA.

As previously reported, the neurons growing in mono-cultures did not develop synchronous activity (Figure S2) whereas the hiPSC-derived co-cultures from all the cell lines developed synchronous network bursting (NB) activity by 5 weeks of differentiation (35DIV, Figure 2A, Figure S3A). Overall, the emergence of the NBs was accompanied by an increase in mean firing rate (MFR, Figure 2B, Figure S3A) that continued throughout neuronal maturation. In addition, all the lines presented sustained network burst frequency (NBF) and network burst duration (NBD) from 5 weeks onward (35DIV, Figure 2C-D, Figure S3A). The NBF and NBD had distinctive patterns between cell lines. After 8 weeks of differentiation, a notable increase in the NBD was observed in one out of the 5 lines (49-56DIV, Figure 2D, Figure S3A). Similarly, the mean inter-spike interval (ISI) within NB decreased gradually after 5 weeks of differentiation in all cell lines, indicating increasing spiking frequency within the NBs (35DIV, Figure 2E, Figure S3A). Taken together, the NGN2-expressing neurons generated sustained synchronous activity in co-cultures with hiPSC-derived astrocytes after 5 weeks of differentiation.

**Figure 2:**
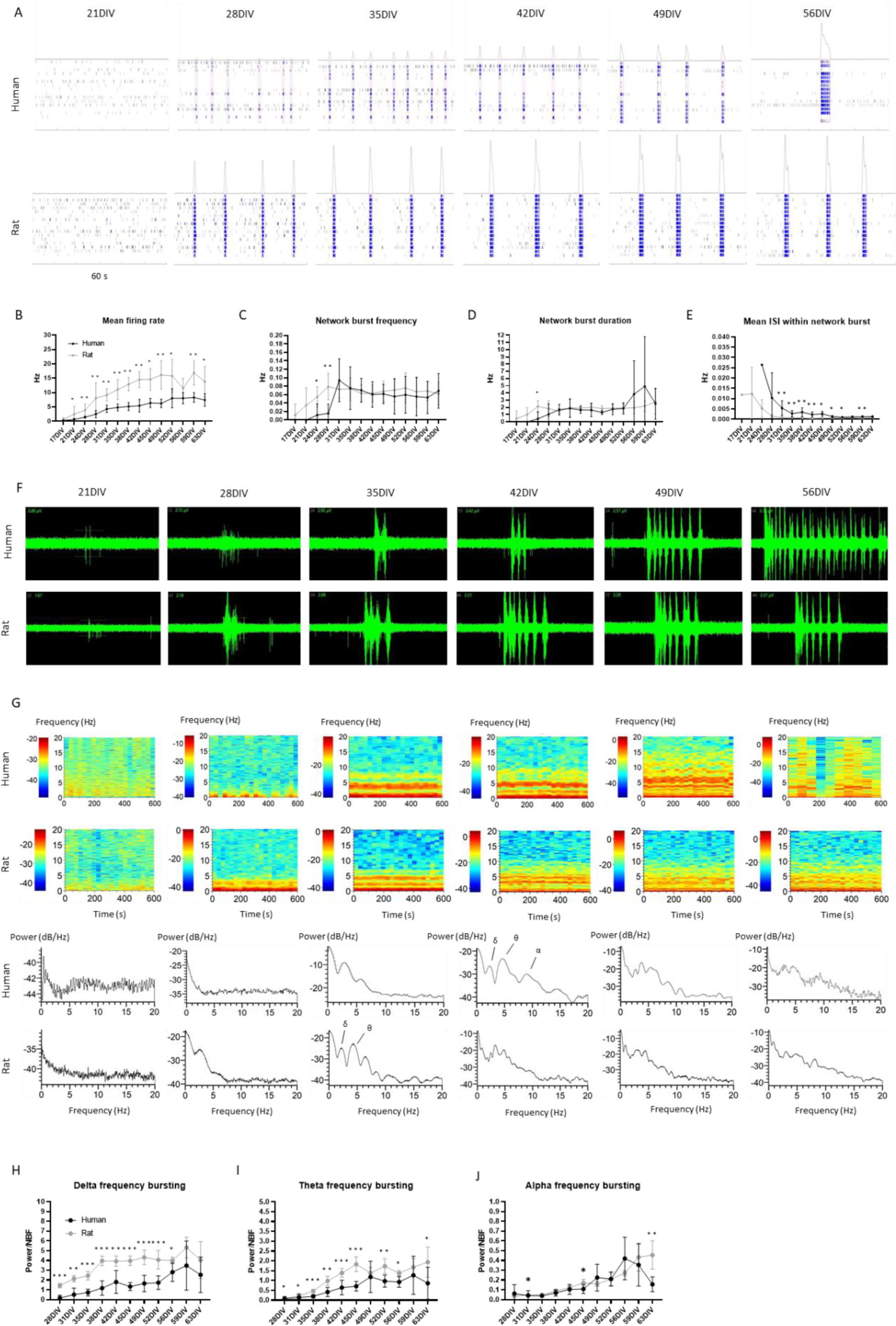
Timeline of the functional development of neurons in co-cultures with hiPSC-derived astrocytes and rat astrocytes. A. Development of network bursting activity in a typical cell line. B. Neuronal MFR increased across time in neuronal co-cultures with human-derived or rat astrocytes. The mixed-species co-cultures displayed higher MFR throughout maturation. C. Development of NBF in hiPSC-derived and mixed-species co-cultures varied between cell lines. D. The NBD is stable until 49DIV but exhibits high variability in later time points (> 56DIV) of hiPSC-derived co-cultures. E. The mean ISI within NB decreased gradually in both culture types. The hiPSC-derived cultures displayed higher mean ISI within NB across differentiation. (n = 5 cell lines, data was collected from 1 experiment) F. Neurons developed high frequency bursting activity within NBs in both hiPSC-derived and mixed-species co-cultures. G. The spectrograms and power spectral densities (PSD) of the raw signal show increase in the high frequency bursting across time. By 5-6 weeks (35 DIV– 42 DIV), distinct activity peaks were distinguishable at delta (δ), theta (θ), and alpha (α) frequencies. H-I. The mixed-species cultures displayed greater bursting activity at delta and theta frequencies than the hiPSC-derived cultures. K. Alpha frequency bursting increased in both culture types around 8 weeks of differentiation (56 DIV). (n= 8 samples from 1 cell line, data was collected from 1 experiment, Mann-Whitney U test was used for the comparisons. * signifies p < 0.05, ** signifies p < 0.01, ns = non-significant)

In concordance with previous studies (2,5), the-mixed-species co-cultures developed synchronous NB activity by 4 weeks of differentiation (28DIV, Figure 2A, Figure S3B) suggesting that neurons in co-cultures with human astrocytes developed slightly slower than in co-cultures with rat astrocytes. The mixed-species co-cultures displayed higher MFR than the fully human cultures throughout differentiation (Figure 2B, Figure S3B) whereas the NBF and NBD converged to similar values in the mixed-species cultures as observed for fully human cultures (Figure 2C-D, Figure S3B). The mean ISI within NB was lower in the mixed-species cultures than in the fully human cultures across maturation, indicating higher spiking frequency within NB (Figure 2E, Figure S3B). This difference likely underlied the higher MFR in the mixed-species cultures. Overall, the hiPSC-derived co-cultures and mixed-species co-cultures produced highly similar NB activity whereas the main differences between the models were found in the timing of the functional switch into NB activity and the spiking frequency within NB affecting the overall MFR.

### NGN2-expressing neurons develop complex network bursting activity in co-cultures with human and rat astrocytes

After observing synchronous NB activity in the co-cultures, we investigated the bursting patterns within the NB in more detail. In both hiPSC-derived and mixed-species co-cultures some neuronal lines developed complex bursting activity characterized by visible sub-bursts within the synchronous events (Figure 2F). Complex bursting that gives rise to oscillatory activity has been detected in long-term neuronal cultures such as brain organoids but only after several months of maturation (19–21). To this end, we wanted to explore whether similar phenomenon could occur in 2D co-cultures and studied the development of bursting at high frequencies (> 0.5 Hz) in one of the cells lines. We first investigated power spectral densities (PSD) and spectrograms of the activity at different time points and noticed increased power below 5 Hz around 5 weeks of differentiation in human co-cultures and around 4 weeks of differentiation in the mixed-species co-cultures (35DIV, 28DIV, Figure 2G). As we continued culturing the cells, the distinct bands diffused to a broader frequency spectrum (35-42DIV Figure 2G). We measured the power of the activity at frequencies 0.5-4 Hz (delta), 4-8 Hz (theta) and 8-12 Hz (alpha) and noticed a steady increase in the activity at all frequencies over time in both culture types (Figure 2H-J). To summarize the findings, the hiPSC-derived neurons produced high frequency bursting activity in co-cultures with hiPSC-derived and rat astrocytes.

### Neuronal response to NMDA receptor blocking differs between human and mixed-species co-cultures

After studying the development of neuronal activity across time, we wanted to understand how different neurotransmitter receptors contributed to the activity in the co-cultures. For this purpose, we prepared hiPSC-derived co-cultures from 5 donors and cultured the cells until 5 weeks of maturation. By this time, all the samples had developed synchronous activity. To first investigate AMPA receptor mediated activity in the cells, we treated them with 10 μM AMPA receptor antagonist NBQX (Figure 3A-B). The treatment significantly reduced the MFR (p = 0.0057) and completely abolished NB activity in all the samples (Figure 3D). Similar abolishment of synchronous bursting has been previously observed in neuronal co-cultures with rodent astrocytes (2).

**Figure 3:**
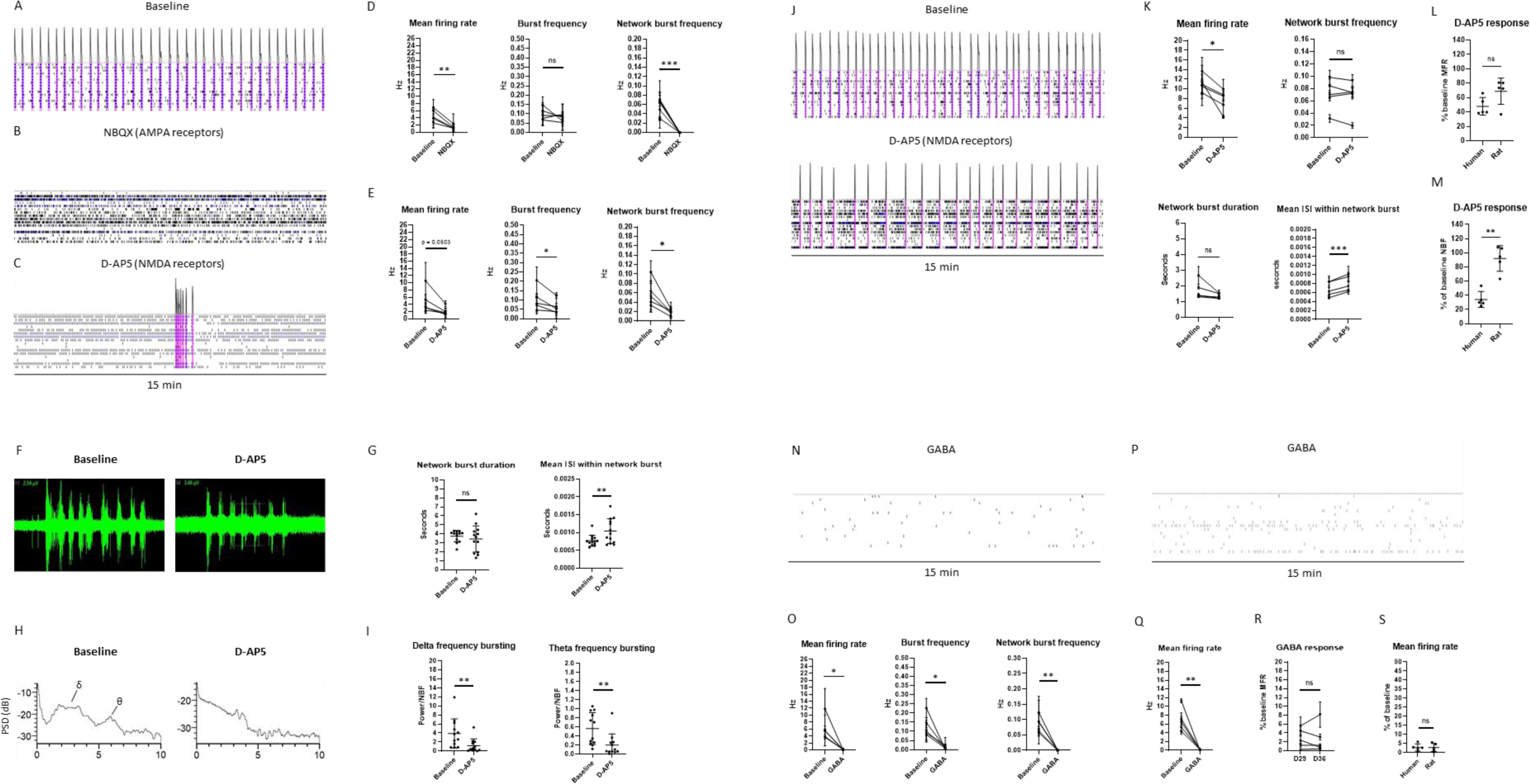
Functional characterization of neuron-astrocyte co-cultures at 5 weeks of differentiation (35DIV). A. Raster plot image showing spiking, bursting and NB activity recorded by each electrode at baseline condition. The hiPSC-derived co-cultures generated regular synchronous activity by 5 weeks of differentiation (35DIV). B. AMPA receptors were blocked with 10 µM NBQX. C. NMDA receptors were blocked with 25 µM D-AP5. D. NBQX induced a significant decrease in MFR and NBF but not burst frequency in the hiPSC-derived cultures. E. D-AP5 induced a decrease in MFR, and a significant decrease in burst frequency and NBF (n = 5 cell lines, data was collected from 2-3 independent experiments.) F. Examples of bursts before and after D-AP5 treatment in hiPSC-derived cultures. G. The NMDA receptor blockage did not affect NBD but significantly increased the mean ISI within network bursts H. PSDs showing changes in neuronal activity at different frequencies after the D-AP5 treatment. I. NMDA receptor blockage significantly reduced bursting at delta and theta frequencies (n= 8-12 samples from one cell line, collected across 3 independent experiments.) J. Raster plot images showing synchronous activity in neuronal co-cultures with rat astrocytes at 5 weeks of differentiation (35DIV), before and after D-AP5 treatment. K. The MFR was significantly reduced after D-AP5 treatment whereas the NBF was not significantly affected by the treatment. The NBD was not affected by the treatment whereas the mean ISI within NB was significantly increased after the D-AP5 treatment. (n = 5 cell lines, data was collected from 1 experiment. Paired t test was used for the statistical comparisons, normal distribution of the data was verified with Kolmogorov-Smirnov test.) L-M. The reduction in NBF after D-AP5 treatment was significantly greater in hiPSC-derived cultures than in the mixed-species cultures at 35DIV. (n = 5 cell lines, data was collected from 1-3 independent experiments. Mann-Whitney U test was used for the comparisons.) N. Response to 100 μM GABA in the hiPSC-derived co-cultures. O. GABA inhibited all bursting and virtually all spiking activity in the hiPSC-derived cultures. P-Q. The mixed-species co-cultures displayed an inhibitory response to GABA at 5 weeks of differentiation. (n = 5 cell lines, data was collected from 2-3 independent experiments for iPSC-derived cultures and 1 experiment for mixed-species cultures. Paired t test was used for the statistical comparisons, normal distribution of the data was verified with Kolmogorov-Smirnov test.) R. The GABA response did not change between 4 and 5 weeks. S. The GABA response did not differ between hiPSC-derived and mixed-species co-cultures. (n = 5 cell lines, data was collected from 1-3 independent experiments. Mann-Whitney U test was used for the comparisons. * signifies p < 0.05, ** signifies p < 0.01, *** signifies p < 0.001, ns = non-significant)

We next investigated the role of NMDA receptors in neuronal activity by treating the cells with 25 μM NMDA receptor antagonist D-AP5 (Figure 3C). The treatment resulted in a nearly significant reduction in the neuronal MFR (p = 0.0503), and a significant reduction in neuronal burst frequency (p=0.0362, Figure 3E). The strongest effect was seen in neuronal NBF that was significantly reduced after the treatment (p=0.00185, Figure 3E). Previous studies have reported that NMDA receptors are necessary for the maintenance of high frequency oscillatory activity in highly mature neuronal cultures (20). Therefore, we also studied the contribution of NMDA receptors to NB patterns in more detail using one of the cell lines. While there was no significant change in the NBD after the D-AP5 treatment, the mean ISI within the NB was significantly increased indicating reduced spiking within the NB (p = 0.0053, Figure 3F-G). We next investigated whether a specific high frequency bursting component within NB was affected by the treatment and measured the effect of D-AP5 on bursting at delta and theta frequencies. As a result, we noticed a reduction in bursting at both frequencies (Figure 3H-I). To summarize the findings, NMDA receptors were identified as critical regulators of neuronal NBF and they were found to enhance high frequency bursting activity within NB across the frequency spectrum.

To study whether the NMDA receptor-mediated activity differed between neuronal co-cultures with human and rat astrocytes, we blocked the NMDA receptors in the mixed-species cultures and compared the results to those obtained with the hiPSC-derived co-cultures (Figure 3J). Given that the neurons matured at different rates between the fully human and mixed-species co-cultures, we studied the NMDA receptor activity at both 4 and 5 weeks in the mixed-species cultures. At 4 weeks (28DIV), the mixed-species cultures displayed a small reduction in MFR and NBF and an increase in the mean ISI within NB after D-AP5 treatment, concordant with the human cultures (Figure S4). After repeating the treatment at 5 weeks, we noticed a reduction in MFR, increase in mean ISI within NB (Figure 3K) but no significant change in NBF (Figure 3K). We compared the reduction in MFR and NBF between the fully human and mixed-species co-cultures at both time points and found that the magnitudes of change in resposens to D-AP5 were overall greater in the hiPSC-derived cultures for MFR and significantly greater for NBF (Figure 3L-M; Figure S4). Thus, the hiPSC-derived cultures displayed greater NMDA receptor activity that strongly affected neuronal NBF. Importantly, this result indicates that the effect of NMDA receptors on NBF was strongly and species-specifically regulated by astrocytes.

Given that astrocytes are known to express functional NMDA receptors, we sought to investigate the effect of neurons and astrocytes on the NMDA receptor function in the co-cultures (22). In general, human astrocytes are larger than rodent astrocytes and could produce more NMDA receptors resulting in stronger NMDA receptor activity (8). Therefore, we first investigated how the number of rat astrocytes in the mixed-species co-cultures affects the NMDA receptor activity. As a result, we found that the astrocyte number did not significantly affect the neuronal response to D-AP5 (Figure S5A). We also quantified the GRIN1 gene expression in the mixed-species and fully human co-cultures but found no evidence of higher GRIN1 expression levels in the hiPSC-derived cultures (Figure S5B). Finally, we conducted ICC staining of GRIN1 to study the distribution of NMDA receptors in neurons and astrocytes. We found that the receptors were concentrated on neuronal somas and diffusely expressed by the astrocytes. The mean GRIN1 intensity did not differ between neurons and astrocytes nor between the different culture conditions. (Figure S5C-D). Overall, these results suggest that the differences in the NMDA receptor function between the hiPSC-derived and mixed-species co-cultures were caused by factors independent of the NMDA receptor expression.

### NGN2-expressing neurons display inhibitory response to GABA

After establishing a central role for the glutamate receptors in maintaining electrophysiological activity, we wanted to investigate the presence of functional GABA receptors in the co-cultures. Although inhibitory neurotransmission is thought to be altered in many developmental and psychiatric disorders, the nature of the GABA response in hiPSC-derived models of the brain has been rarely characterized (12,21,23). We first applied 100 μM GABA to hiPSC-derived co-cultures at 5 weeks of differentiation (Figure 3N) and found that the neurons exhibited a strong inhibitory response to GABA. Altogether, GABA blocked 94-100% of the neuronal spiking activity and consequently all NB activity (Figure 3O). To find out whether the inhibitory response to GABA was a unique phenomenon in the hiPSC-derived cultures, we also investigated GABA responses in mixed-species co-cultures after 4 and 5 weeks of maturation (Figure 3P). Already after 4 weeks, we observed a strong inhibitory GABA response (Figure 3P) that did not change between 4 and 5 weeks (Figure 3Q-R). The effect of GABA on the MFR did not differ between the fully human and mixed-species cultures (Figure 3S). Hence, both human-based and mixed-species co-cultures developed a fully inhibitory response to GABA by 4-5 weeks of differentiation.

### hiPSC-derived co-cultures from patients with schizophrenia show network bursting alterations

We next investigated the suitability of the hiPSC-derived neuron-astrocyte co-cultures for modeling cellular functions underlying psychiatric disorders. We have previously performed transcriptomic profiling for neurons and astrocytes derived from monozygotic twins discordant for TRS, and showed alterations in pathways such as glutamate signaling and synaptic functions in schizophrenia. Using calcium signaling measurements, we have also shown that neuronal NMDA receptor-mediated glutamate response differs between neurons from the affected twins (AT) and neurons from their unaffected co-twins in co-cultures with astrocytes (9,12). To first investigate whether co-cultures derived from these same cell lines display characteristic alterations in network bursting, we differentiated donor-matched neuron-astrocyte co-cultures from hiPSCs derived from 3 monozygotic twin pairs discordant for TRS and 3 unrelated controls. After ICC staining and characterization of the cultures, we found no differences in the number of MAP2 or CUX1 positive neurons and detected no PAX6 and PRPH-positive cells in the co-cultures. The number of S100β and GFAP expressing astrocytes was also similar between the conditions concordantly with our previous results (9) (Figure S6). Full characterization of the astrocytes including ICC staining, qPCR and glutamate uptake assay has been performed in our earlier study revealing no differences between the affected and unaffected astrocytes (9).

We next prepared donor-matched co-cultures on MEA and monitored the development of neuronal activity until 42DIV when all the lines had developed robust NB activity (Figure 4A-B). At this stage, we noticed that both the UT and CTR cultures displayed higher MFR than the AT cultures (Figure 4B, C, Table S2, Figure S7). Overall, we found a prominent donor effect for MFR with 38.77% of variance explained by the donor whereas a similar 38.43% effect was found to be explained by the schizophrenia status. Belonging to the same twin pair did not contribute to the variance (Table S2).

**Figure 4:**
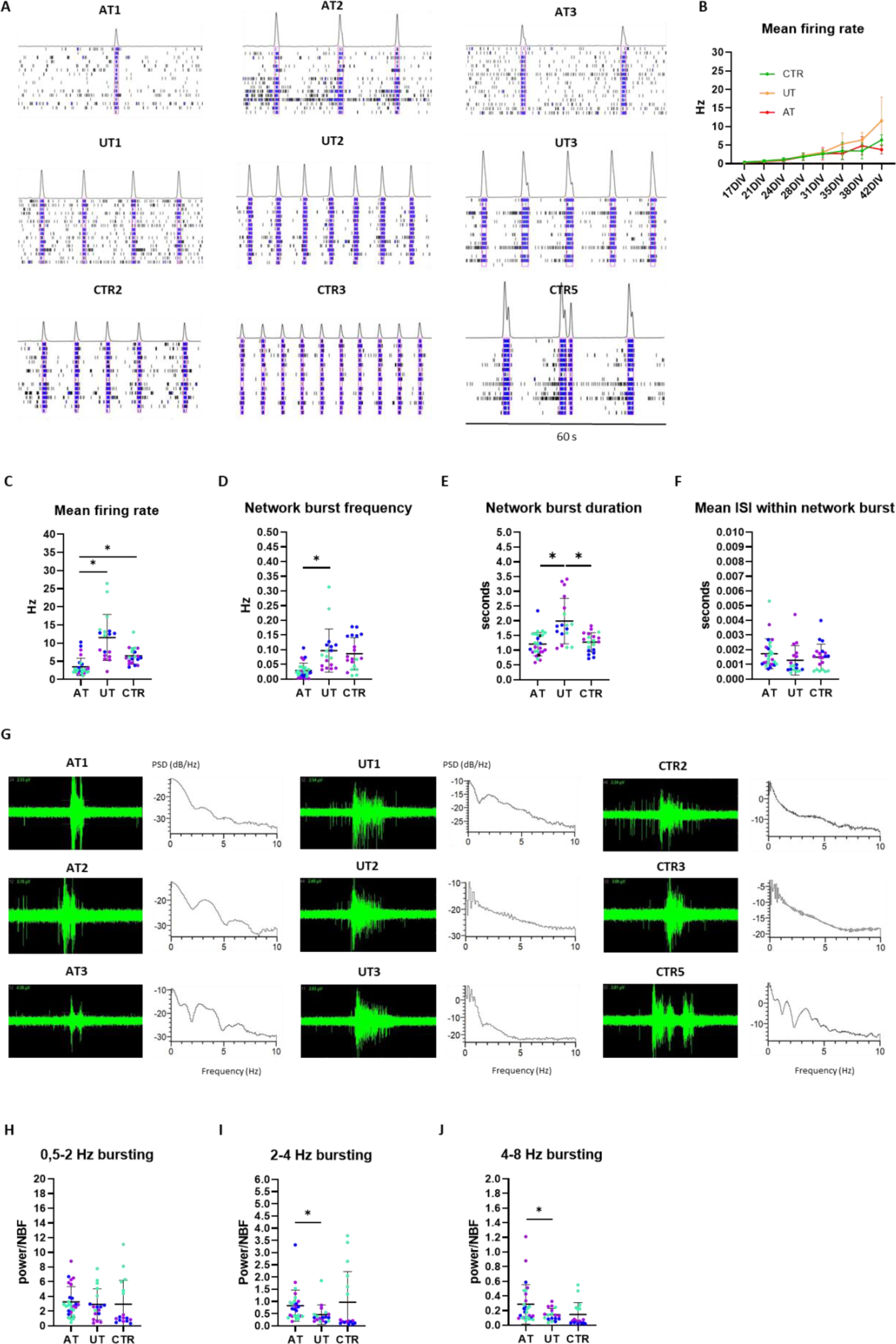
Characterization of neuronal activity in co-cultures derived from monozygotic twins discordant for treatment-resistant schizophrenia and controls. A. Raster plot images showing NB activity in co-cultures derived from affected twins (AT), unaffected twins (UT) and control individuals (CTR) at 42DIV. B. Development of MFR in cultures derived from AT, UT and CTR individuals. C. Co-cultures derived from UT and CTR individuals displayed nominally significantly higher MFR compared to cultures derived from the AT at 42 DIV. D. Cultures derived from UT exhibited nominally significantly higher NBF compared to cultures derived from AT. E. Cultures derived from UT exhibited nominally significantly higher NBD compared to the cultures derived from AT and CTR. F. The mean ISI within NB did not differ between the cultures. G. Raw signal showing burst shapes in cultures derived from AT, UT and CTR, and PSDs showing the distribution of frequency components in the donor-specific cultures at 42 DIV. H-J. Analysis of high-frequency bursting in AT, UT and CTR cultures revealed differences in 2-4 Hz bursting and 4-8 Hz bursting between AT and UT cultures. (n = 3 cell lines per group, data was collected from 1-2 independent experiments. P-values were derived from ANOVA using a general mixed linear regression model, the figures display uncorrected p-values, * signifies p < 0.05, ns = non-significant, the colors of the data points represent results from different cell lines.)

We further scrutinized the underlying factors of the differences in MFR and found that the UT cultures displayed nominally significantly higher NBF (p = 0.02389, p_adj_ = 0.13473) and NBD (p = 0.02236, p_adj_ = 0.13473) than the AT cultures (Figure 4D). In addition, the UT cultures exhibited higher NBD than the CTR cultures (p = 0.03649, p_adj_ = 0.13473, Figure 4E). Although we found no difference in the mean ISI within NB between the groups (Figure 4F), we did detect differences in the burst shapes between the AT and UT cultures. At this stage, the AT cultures displayed more distinct sub-bursts within the NB (Figure4G). After quantifying neuronal bursting activity at high frequencies, we found stronger activity at 2-4 Hz and 4-8 Hz in the AT cultures compared to the UT cultures (Figure 4J). In conclusion, using human-based co-cultures, we were able to show differences in the activity of neurons from patients with schizophrenia and unaffected controls originating from NB patterns. Here, the difference in the activity between AT and UT cultures resulted from higher frequency and longer duration of NBs in the UT cultures.

### The role of neuronal and astrocyte donor on NBF and NBD

To investigate the role of neurons and astrocytes in regulating NBF and NBD, we selected co-cultures derived from one twin pair for more detailed characterization. We prepared mixed-donor cultures containing neurons and astrocytes from the UT (NU+AU), neurons from the UT and astrocytes from the AT (NU+AA), neurons and astrocytes from the AT (NA+AA) and neurons from the AT and astrocytes from the UT (NA+AU, Figure 5A). We measured the neuronal activity at 6 weeks and compared the cultures in terms of NBF and NBD (Figure 5B-C). Interestingly, when the UT neurons were cultured with AT astrocytes, their NBF was significantly reduced (p = 0.0098). Moreover, when AT neurons were cultured with UT astrocytes, their NBF was significantly increased (p < 0.0001). This finding suggested that astrocytes critically influenced the neuronal NBF. When we measured the NBD in the different cultures, we found that the NBD was slightly higher in cultures containing UT neurons than in cultures containing AT neurons suggesting that neurons, instead of astrocytes, regulated the NBD (Figure 5C). Taken together, we found that NBF and NBD were differentially influenced by neurons and astrocytes, and that NBF was strongly affected by the astrocyte donor.

**Figure 5:**
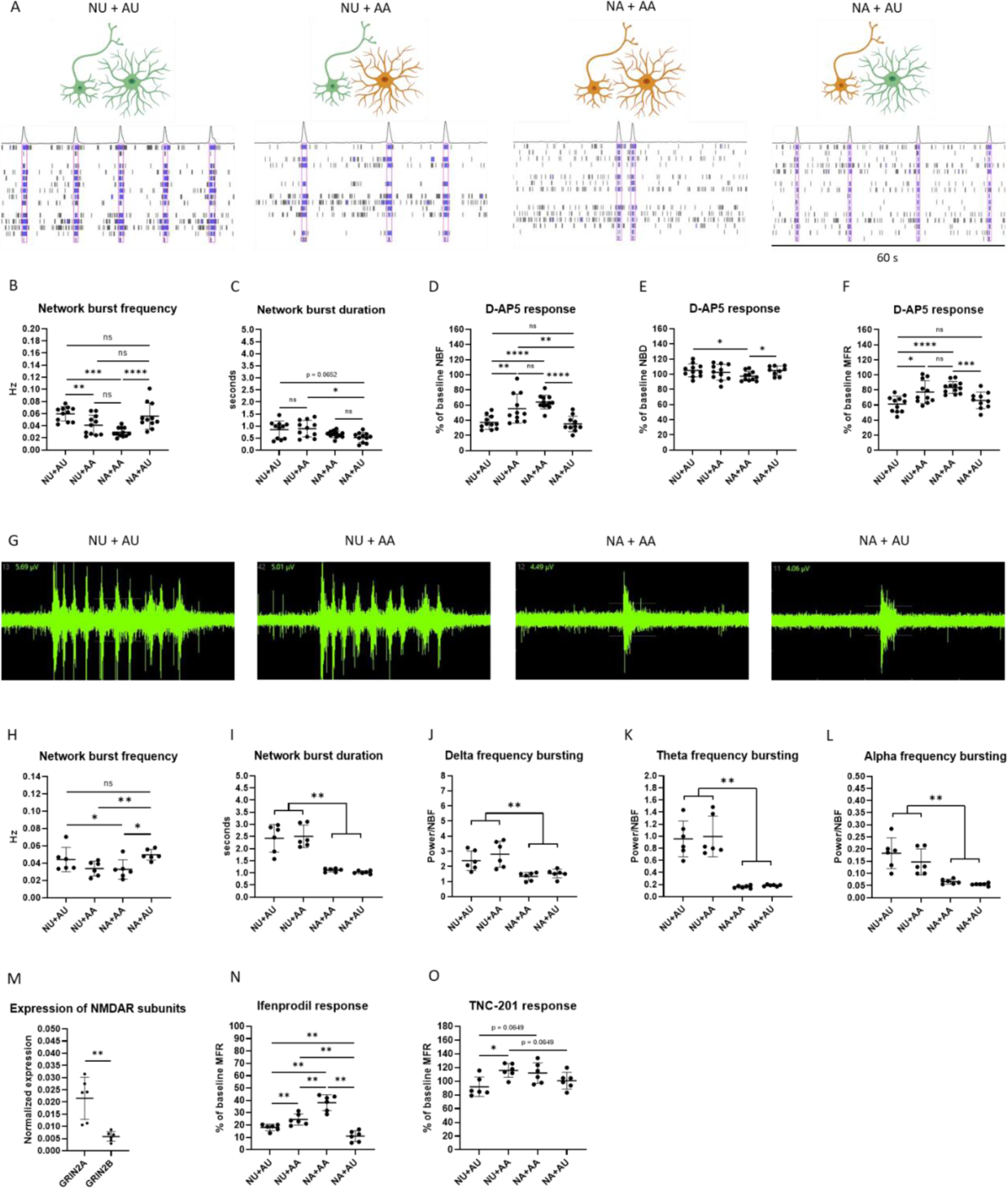
Characterization of neuronal activity in mixed-donor co-cultures derived from one patient affected with schizophrenia and an unaffected twin. A. Examples of NB activity in cultures with neurons and astrocytes from UT (NU+AU), neurons from UT and astrocytes from AT (NU+AA), neurons and astrocytes from AT (NA+AA), and neurons from AT and astrocytes from UT (NA+AU). B. NU+AU cultures displayed significantly higher NBF than NA+AA cultures. NU+AA cultures displayed a significantly lower NBF than NU+AU cultures and NA+AU displayed significantly higher NBF than NA+AA cultures. C. The NBD was slightly higher in cultures containing UA than cultures containing AA. D. The effect of D-AP5 on NBF was significantly stronger in NU+AU cultures compared to NA+AA cultures. The NU+AA cultures displayed a significantly weaker D-AP5 effect of NBF compared to NU+AU cultures. The effect of D-AP5 on NBF was similar to the level of NU+AU cultures when NA were cultured with AU. E. The effect of D-AP5 on NBD was independent of neuron or astrocyte donor. F. The effect of D-AP5 on MFR reflected the effect of D-AP5 on NBF. (n = 10-12 samples collected across 2 independent experiments. Mann Whitney U test was used for the statistical comparisons. **** signifies p < 0.0001, *** signifies p < 0.001, ** signifies p < 0.01, * signifies p < 0.05, ns = non-significant) G. Raw signal showing bursts in mixed-donor cultures at 8 weeks of differentiation. H. The NBF was significantly lower in cultures containing AA compared with cultures containing AU. I. NBD was significantly elevated in cultures containing NU compared with cultures containing NA. J-L. The power of delta, theta and alpha frequency bursting was higher in cultures containing NU compared to cultures containing NA. M. Expression of GRIN2A and GRIN2B subunits in control co-cultures at 5 weeks. N. The cultures containing AU displayed the highest response to GLUN2B-specific blocker Ifenprodil (10 μM). O. GLUN2A-specific blocker TNC-201 (3 μM) slightly increased activity in cultures containing AA. (n = 6 samples per condition, data was collected from 1 experiment. Mann Whitney U test was used for the statistical comparisons, ** signifies p < 0.01, * signifies p < 0.05, ns = non-significant)

Given that we earlier evidenced the role of astrocytes in regulating NBF via NMDA receptors, we wanted to investigate whether the differences in NBF in cultures from affected and unaffected individuals were also mediated through NMDA receptors. We exposed the cultures to D-AP5 and measured the effect of the treatment on NBF. We found that the NBF in NU+AU cultures was reduced significantly more than in the NA+AA cultures (p < 0.0001, Figure 5D), confirming that the UT cultures expressed stronger NMDA receptor-mediated activity. We also found that the NMDA receptor-mediated effect on NBF in AT neurons was fully rescued when cultured with UT astrocytes. Likewise, the NMDA receptor-mediated effect on NBF was significantly attenuated in UT neurons when cultured with AT astrocytes (p = 0.0066, Figure 5D). Hence, the astrocytes had donor-specific, NMDA receptor-dependent effects on neuronal NBF. We also studied the effect of D-AP5 on NBD and found minor changes that were not explained by the neuron or astrocyte donor (Figure 5E). When we measured the change in MFR between the groups after NMDA receptor blockage, we observed a pattern of higher response in cultures containing UT astrocytes compared with cultures containing AT astrocytes (Figure 5F). This pattern reflected that of D-AP5 effect on NBF indicating that the NMDA receptor mediated activity primarily affected the NBF in the co-cultures. To test whether the phenomenon was consistent, we also prepared co-cultures using cells from the other 2 discordant twin pairs, and measured the effect of D-AP5 on MFR and NBF. In case of both pairs, the UT cultures displayed a significantly higher D-AP5 response compared to the AT cultures strengthening the idea that attenuated NMDA receptor activity was associated with schizophrenia (Figure S8).

We next wanted to investigate the effect of the neuron and astrocyte donor on the activity patterns within NB. For this purpose, we continued culturing the cells until 8 weeks to see oscillatory activity within the network events (Figure 5G). At this stage, we observed higher NBF and significantly greater NBD in cultures with UT neurons compared to cultures with AT neurons (p = 0.0022, Figure 5H-I). The oscillatory bursting at delta, theta and alpha frequencies was also significantly higher in cultures with UT neurons compared to cultures with AT neurons (p = 0.0022, Figure 5J-L). Notably, the cultures with neurons from the same donor displayed highly similar NBD and high frequency bursting patterns suggesting that these features were exclusively regulated by neurons. The prolonged culturing of the cells also revealed that the MFR and NBD deviated progressively between cultures containing neurons from a different donor (Figure S9). This result suggested that the differences observed in neuronal activity at early stages were sustained across development.

Finally, we wanted to elucidate the role of NMDA receptors containing either GLUN2A or GLUN2B subunits in the electrophysiological activity of the co-cultures. The two paralogs have different temporal expression profiles during brain development with GRIN2B, which encodes the GluN2B subunit of NMDA receptors, being highly expressed prenatally and GRIN2A, which encodes the GluN2A subunit of NMDA receptors, becoming the dominant subtype after birth (16). In our co-cultures, we observed high GRIN2A mRNA levels already after 5 weeks of differentiation suggesting their potential contribution to the NMDA receptor-mediated activity (Figure 5M). Furthermore, gene disabling variants in *GRIN2A* were recently found to be associated with schizophrenia (24). We first used a GLUN2B-specific blocker Ifenprodil (10 μM) to block the NMDA receptors in the mixed-donor co-cultures (Figure 5N) and detected a strong response in all cultures. The most prominent response was observed in cultures containing UT astrocytes suggesting that the differences in NMDA receptor function between the cultures were regulated by astrocytes through GLUN2B-containing receptors. We next blocked the GRIN2A-containing receptors with TCN-201 (3 μM) and observed a small increase in the MFR in cultures with AT astrocytes (Figure 5O). Based on this result, we concluded that GLUN2A-containing receptors played at most a minor role in regulating neuronal activity at this stage.

## Discussion

During the past few years, the importance of astrocytes in regulating neuronal activity has been increasingly recognized. Several hiPSC-based studies have demonstrated how the presence of astrocytes is required for the development of complex neuronal network-level activity *in vitro* (2,6,19,25). Astrocytes are also known to play a role in the pathogenesis of conditions such as schizophrenia and Alzheimer’s disease (4,9,10). Hence, there is an increasing need for the use of donor-specific astrocytes in hiPSC-based models of the brain.

Here, we established fully human neuron-astrocyte co-cultures derived from hiPSCs. Using MEA, we detected development of synchronous NB activity after 5 weeks of neuronal differentiation in these cultures. Previous reports have shown that NGN2-expressing neurons develop synchronous activity with rodent astrocytes by 4 weeks of differentiation (2,5). Hence, the hiPSC-derived cultures displayed a one-week delay in their development in comparison to mixed-species cultures. However, overall the hiPSC-derived and mixed-species cultures displayed highly similar activity patterns across development.

In addition to NB activity, we detected complex high frequency bursting activity developing within the synchronous events in both fully human and mixed-species cultures. Previous studies have reported that NGN2-expressing neurons do not develop oscillations corresponding to those observed in preterm human infants (20). Interestingly, the development of oscillatory activity has been observed in different types of iPSC-based models of the brain around the time when astrocytes start differentiating, typically after several months of maturation (19,20,25). Furthermore, boosting astrocyte differentiation in these cultures has been shown to propone the development of oscillatory activity (25). Therefore, using co-cultures of neurons and astrocytes could be the key to generating models with mature neuronal activity within a short period of time.

We performed pharmacological characterization of the co-cultures to study the contribution of different receptor types on the generation of neuronal activity. Pharmacological blockage of NMDA receptors induced a strong reduction in NBF in the hiPSC-derived co-cultures in contrast to the mixed-species co-cultures. Similar, minor responses to NMDA receptor blockage have also been reported in other studies using NGN2-expressing neurons cultured with rodent astrocytes (2,26,27). This finding not only indicates that astrocytes influence the generation of synchronous activity in neuronal networks through NMDA receptors but also that this process is differently regulated by human and rodent astrocytes. Although we were not able to unravel the mechanisms of this regulation, it is known that glutamate released from astrocytes can bind to extra-synaptic NMDA receptors in neurons, resulting in their synchronous excitation (28). This suggests that the differences observed between the human and rat astrocytes could originate from differences in glutamate release and uptake. In addition, soluble factors released by astrocytes have been shown to specifically activate synaptic GLUN2B-containing NMDA receptors in neurons (29). On the other hand, astrocytes are also known to express functional NMDA receptors that regulate astrocyte calcium influx (22). In general, the development of hiPSC-derived neurons and astrocytes *in vitro* follows a similar time course to human brain development (30). Thereby, both iPSC-derived astrocytes and rat astrocytes (E18) used in this study represent a developmental stage corresponding to late pregnancy.

Given that we had observed signs of highly developed activity in our cultures, we also wanted to examine the effect of GABA on our neurons. Typically, GABA is thought to have an excitatory effect on young, prenatal neurons, and after birth, the effect switches to inhibition (31). Thereby, the excitatory-inhibitory switch in neurons is associated with the maturational stage of the cells. A few studies have reported such a switch occurring in hiPSC-derived models of the brain (6,25,32). Notably, we were able to show that our neurons displayed a fully inhibitory response to GABA by 4- 5 weeks of differentiation when cultured with human or rodent astrocytes. This finding shows that hiPSC-derived neurons can develop a mature, inhibitory response to GABA in a short period of time and in the absence of inhibitory neurons.

Finally, we used the co-culture model to investigate differences between brain cells derived from patients with schizophrenia and unaffected individuals. We were able to detect differences in the NB patterns between healthy and affected co-cultures at 6 weeks of differentiation. Previously, only one study has characterized neuronal NB patterns using hiPSC-derived neurons from patients with schizophrenia (33). This article also found evidence for reduced neuronal activity and NBF in the affected neurons after 6 weeks of differentiation. Along with our results, this finding indicates that the presence of NB activity may be critical in order to detect disease-related functional alterations in neurons. We were able to demonstrate how the NBF was regulated through GRIN2B-containing NMDA receptors. Despite the small sample size, our findings are in line with earlier results showing altered NMDA receptor dynamics in clozapine-responsive schizophrenia (34).

In addition to NBF, we also detected differences in NBD and high frequency bursting activity within NB between neurons from the AT and UT. These differences were not associated with NMDA receptor function. In a recent hiPSC-based study, altered Na^+^ channel function and subsequent decrease in neuronal excitability were found to correlate with the severity of positive symptoms in the donor patients with schizophrenia (23). Given that the activity within NB was largely regulated by neurons in the co-cultures, changes in intrinsic neuronal excitability could explain the observed difference. In support of this hypothesis, mutations in *SCN1A* gene coding for Na_v_1.1 subunit were recently linked to a bursting phenotype with distinct sub-burst in patient-derived neurons (35). Overall, there seem to be multiple components to neuronal dysfunction in schizophrenia. As the altered NMDA receptor function has been associated specifically with TRS (34), the decreased neuronal excitability appears to be a more general hallmark of the disorder (23).

Intriguingly, we have consistently found differences between iPSC-derived brain cells from identical twins that represent the same genetic background (9,12). In addition, other studies using hiPSCs from monozygotic twins discordant for schizophrenia have found differences in neuronal activity and gene expression patterns between the AT and UT cultures (36–39). These differences could be explained by *de novo* mutations carried by either one of the twins. Indeed, we have earlier detected several non-overlapping copy number variants between our twin lines (12). To further illuminate the disease mechanisms of schizophrenia in monozygotic twins, studies focusing on genetic and epigenetic differences between the twins would be needed.

## Acknowledgements

This work was supported by Academy of Finland (grant 334525, JK) and Sigrid Juselius Foundation (JK).

## Author contibutions

Conceptualization, J.K., O.P., J.T., N.R.; Methodology, J.K., O.P., J.T., N.R.; Investigation, N.R., M.K., N.J., N.K., I.W.; Writing – Original Draft, N.R.; Writing – Review & Editing, N.R., Š.L., O.P.; Visualization, N.R.; Formal Analysis, N.R., O.P.; Funding Acquisition, J.K.

## Disclosures

The authors declare no competing interests.

## Notes

### Competing Interest Statement

The authors have declared no competing interest.

