## Supplemental tables and figures for "Astrocytes regulate neuronal network burst frequency through NMDA receptors species- and donor-specifically"

**Supplemental Table 1:** *List of hiPSC lines used in this study.*

| Cell line | Status | Age at biopsy | Sex | Medication |
|---|---|---|---|---|
| CTR1 | control | 44 | Male | - |
| CTR2 | control | 49 | Female | - |
| CTR3 | control | 64 | Female | - |
| CTR4 | control | 63 | Male | - |
| CTR5 | control | 50 | Female | - |
| AT1 | affected twin | 47 | Female | clozapine |
| AT2 | affected twin | 69 | Female | Previously clozapine, now sertindole and quetiapine |
| AT3 | affected twin | 45 | Female | clozapine |
| UT1 | unaffected twin | 47 | Female | - |
| UT2 | unaffected twin | 69 | Female | - |
| UT3 | unaffected twin | 45 | Female | - |

**Supplemental Table 2:** *Statistical comparison of electrophysiological results for twins discordant for schizophrenia and unrelated controls at 42 DIV.*

| Variable | AT vs CTR | AT vs UT | UT vs CTR | Proportion of variance (ID/PairID/Status/Residual) |
|---|---|---|---|---|
| MFR | -2.818<br>(1.060)<br>p=0.04491<br>$p_{adj}$ = 0.13473 | -8.043<br>(2.887)<br>p=0.02582<br>$p_{adj}$ = 0.13473 | 5.297<br>(2.908)<br>p=0.1034<br>$p_{adj}$ = 0.24127 | 38.77/0/38.43/22.80 |
| NBF | -0.05774<br>(0.03010)<br>p=0.09087<br>$p_{adj}$ = 0.23853 | -0.07025<br>(0.02333)<br>p=0.02389<br>$p_{adj}$ = 0.13473 | 0.01283<br>(0.03924)<br>p=0.7446<br>$p_{adj}$ = 0.93042 | 50.26/0/15.93/33.81 |
| NBD | -0.03953<br>(0.20347)<br>p=0.8464<br>$p_{adj}$ = 0.96090 | -0.7559<br>(0.2574)<br>p=0.02236<br>$p_{adj}$ = 0.13473 | 0.7082<br>(0.2789)<br>p=0.03649<br>$p_{adj}$ = 0.13473 | 26.31/0/30.21/43.48 |
| mean ISI within NB | 0.0001955<br>(0.0004065)<br>p=0.6365<br>$p_{adj}$ = 0.93042 | 0.0004576<br>(0.0003919)<br>p=0.28<br>$p_{adj}$ = 0.53455 | -0.0002896<br>(0.0005882)<br>p=0.6255<br>$p_{adj}$ = 0.93042 | 32.47/0/0/67.53 |
| 0.5-2 Hz bursting | 0.5457<br>(1.7296)<br>p=0.7532<br>$p_{adj}$ = 0.93042 | 0.4246<br>(1.1635)<br>p=0.7165<br>$p_{adj}$ = 0.93042 | 0.1237<br>(1.8729)<br>p=0.9474<br>$p_{adj}$ = 0.96090 | 62.80/0/0/37.20 |
| 2-4 Hz bursting | -0.05501<br>(0.62841)<br>p=0.9303<br>$p_{adj}$ = 0.96090 | 0.3674<br>(0.1656)<br>p=0.03211<br>$p_{adj}$ = 0.13473 | -0.4776<br>(0.6300)<br>p=0.459<br>$p_{adj}$ = 0.80325 | 58.99/0/0/41.01 |
| 4-8 Hz bursting | 0.14544<br>(0.09605)<br>p=0.1604<br>$p_{adj}$ = 0.33684 | 0.1475<br>(0.06163)<br>p=0.04062<br>$p_{adj}$ = 0.13473 | -0.004097<br>(0.083547)<br>p=0.9609<br>$p_{adj}$ = 0.96090 | 21.72/0/5.3/72.97 |

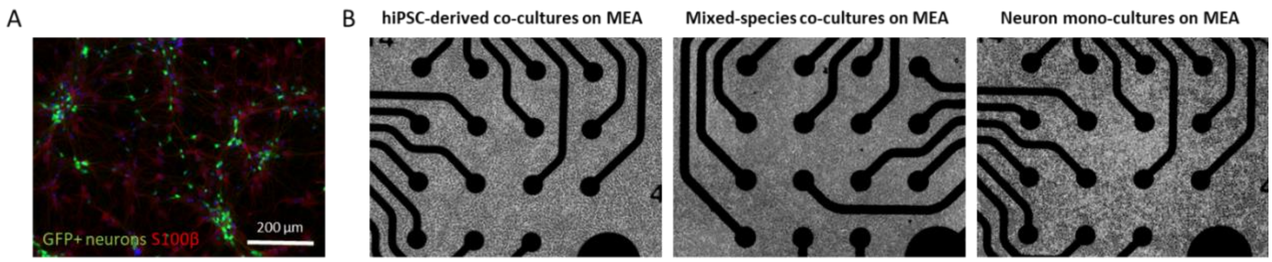


**Supplemental Figure 1**: hiPSC-derived neurons and astrocytes in co-cultures. A. Neurons were labeled with GFP for ICC characterization. B. Co-cultures and neuronal mono-cultures plated on MEA.

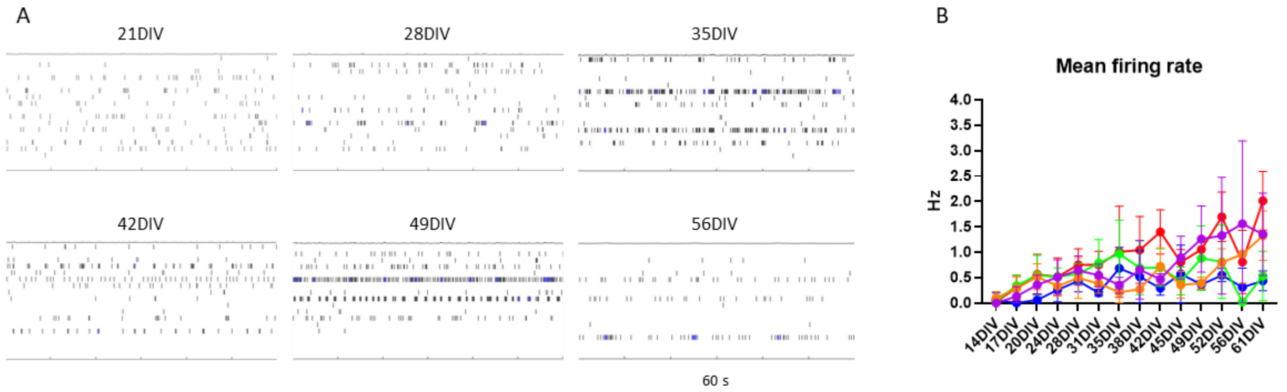


**Supplemental Figure 2**: Functional characterization on neuronal mono-cultures on MEA. A. Raster plot images showing neuronal spiking and bursting activity. B. MFR across time measured from 5 cell lines.

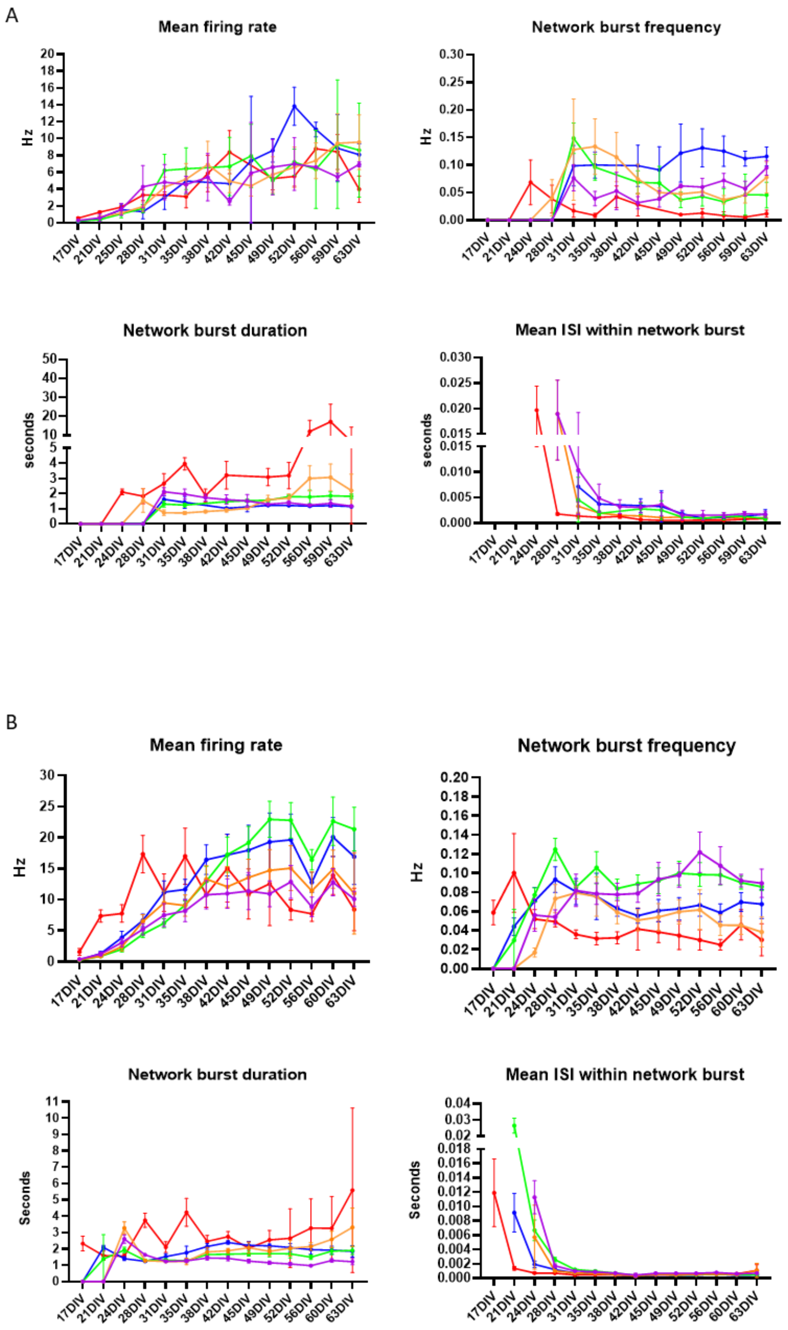


**Supplemental Figure 3**: Timeline of the functional development of neurons with hiPSC-derived astrocytes (A) and rat astrocytes (B).

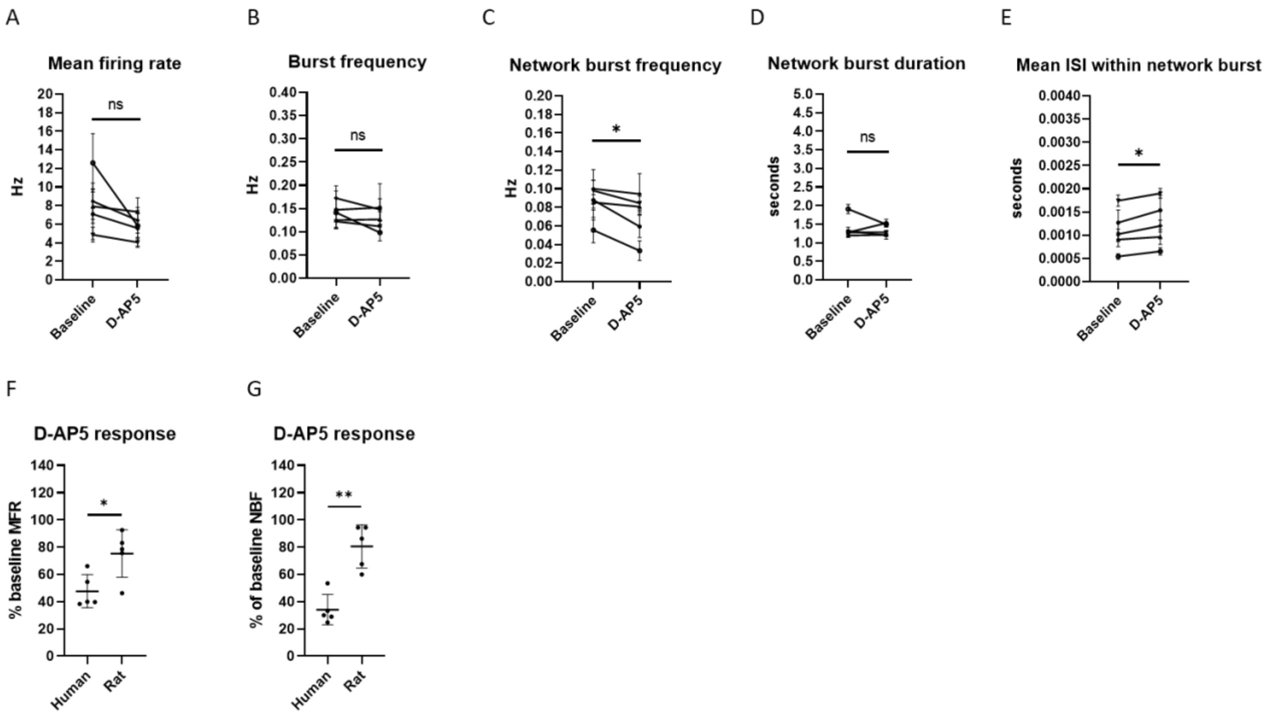


**Supplemental Figure 4:** Characterization of neuronal D-AP5 response in co-cultures with rat astrocytes at 4 weeks of differentiation. A-E. The NMDA receptor blockage had a significant effect on NBF and mean ISI within NB in the mixed-species cultures. F-G. The hiPSC-derived cultures displayed a stronger response to D-AP5 than the mixed-species co-cultures. (n = 5 cell lines, data was collected from 1 experiment. For A-E: Paired t test was used for the statistical comparisons, normal distribution of the data was verified with Kolmogorov-Smirnov test. For F-G: Mann-Whitney U test was used for the statistical comparisons. * signifies $p < 0.05$, ** signifies $p < 0.01$, ns = non-significant)

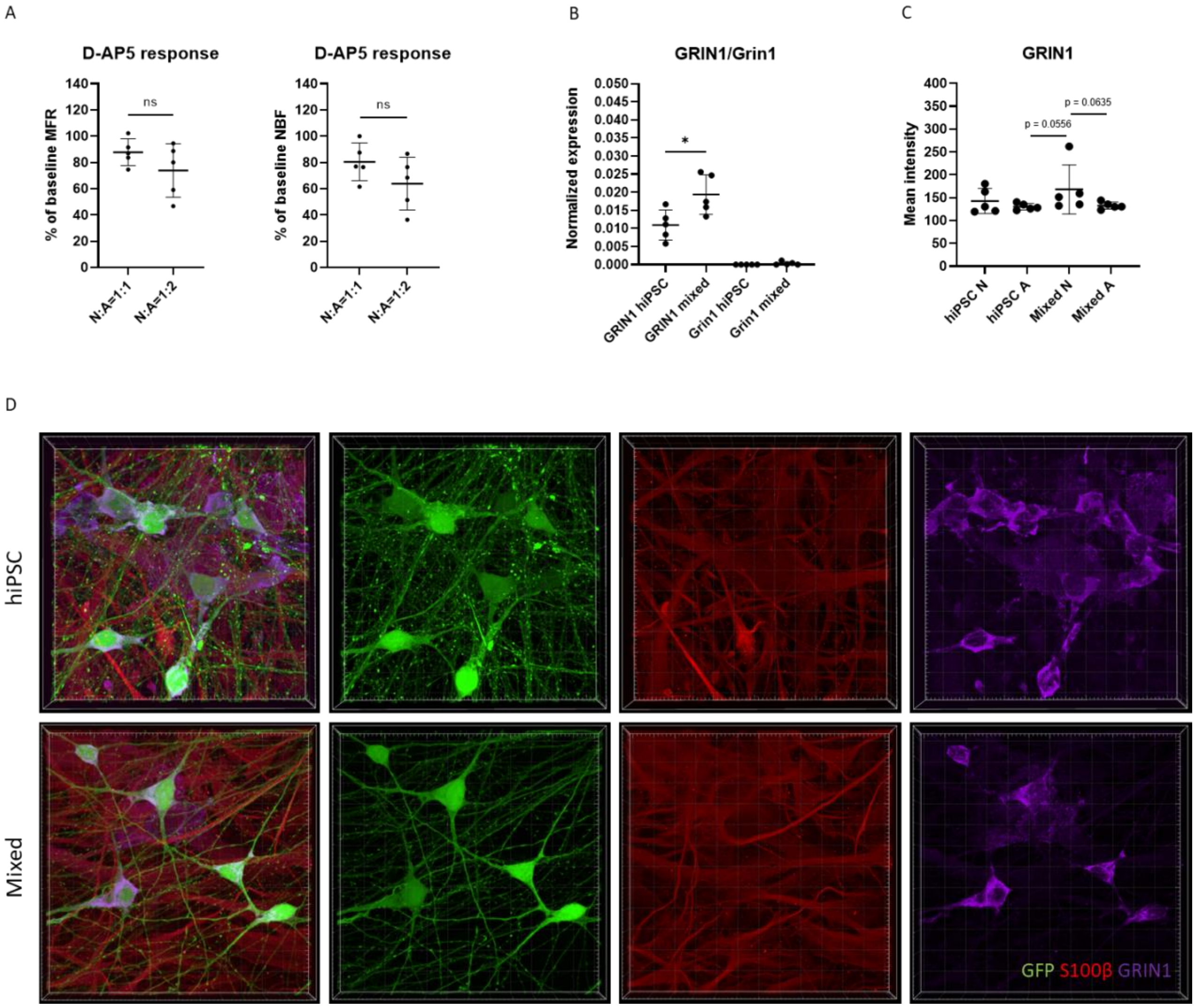


**Supplemental Figure 5**: Characterization of NMDA receptors in neuron-astrocyte co-cultures. A. The number of astrocytes did not significantly affect the NMDA receptor activity in mixed-species co-cultures. B. qPCR quantification of human-specific GRIN1 and rat-specific Grin1 subunits in hiPSC-derived and mixed-species co-cultures. C-D. ICC quantification of GRIN1 subunit expression in hiPSC-derived and mixed-species co-cultures revealed no differences in NMDA receptor expression between cell types and culture conditions. (n=5 cell lines, data was collected from 1 experiment, Mann-Whitney U test was used for the statistical comparisons, * signifies $p < 0.05$, ns = non-significant)

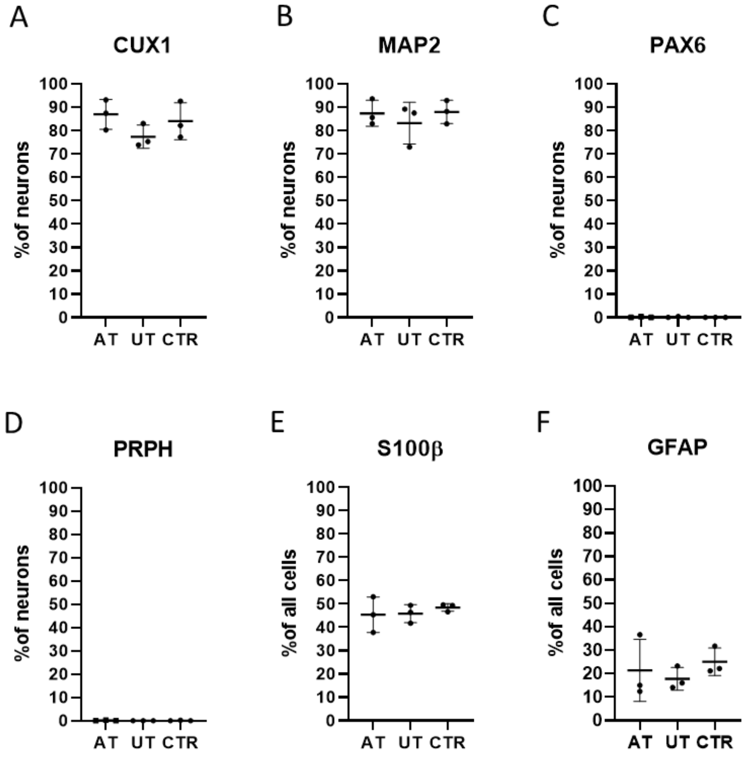


**Supplemental Figure 6:** Characterization of neuron-astrocyte co-cultures derived from monozygotic twins discordant for treatment-resistant schizophrenia. A-B. Neurons in cultures derived from affected twin (AT), unaffected twin (UT) and control cultures (CTR) were CUX1 and MAP2 positive. C-D. PAX6 and PRPH expressing cells were not detected in the cultures. E-F. Approximately 50% of the cells in all co-culture conditions expressed astroglial marker S100β and on average 20 % of the cells expressed GFAP. (n = 3 cell lines per group, data was collected from 2 independent experiments, each data point represents 2-4 replicate samples)

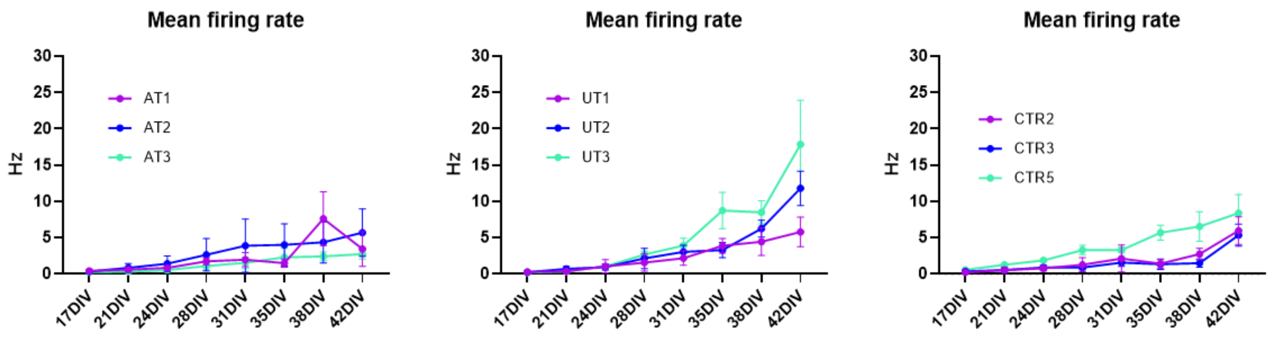


**Supplemental Figure 7:** MFR measured from neuron-astrocyte co-cultures from affected twins (AT), unaffected twins (UT) and control individuals (CTR).

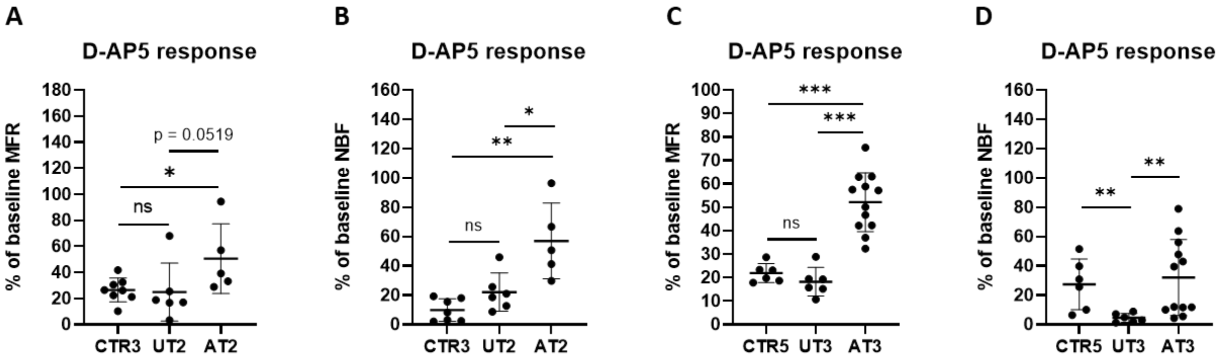


**Supplemental Figure 8:** D-AP5 responses in twin pairs 2 (A-B) and 3 (C-D). The UT cultures displayed a stronger response to D-AP5 in terms of MFR and NBF compared to AT cultures. (n = 5-12 samples per condition. The data was collected across 1-2 independent experiments)

A

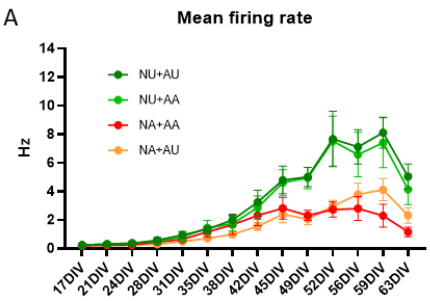


| | NU+AU vs NU+AA | NU+AA vs NA+AA | NA+AA vs NA+AU | NU+AU vs AA+NU | NU+AU vs NA+AA | NU+AA vs NA+AU |
|---|---|---|---|---|---|---|
| 17DIV | ns | ns | ns | ns | ns | ns |
| 21DIV | ns | ns | ns | ns | ns | ns |
| 24DIV | ns | p = 0.0411 | ns | ns | ns | p = 0.0087 |
| 28DIV | ns | ns | p = 0.0411 | p = 0.0022 | ns | p = 0.0260 |
| 31DIV | ns | ns | ns | p = 0.0022 | p = 0.0260 | p = 0.0022 |
| 35DIV | ns | ns | p = 0.0260 | p = 0.0022 | ns | p = 0.0022 |
| 38DIV | ns | ns | p = 0.0260 | p = 0.0022 | ns | p = 0.0260 |
| 42DIV | ns | ns | p = 0.0411 | p = 0.0022 | ns | p = 0.0022 |
| 45DIV | ns | p = 0.0152 | ns | p = 0.0022 | p = 0.0043 | p = 0.0043 |
| 49DIV | ns | p = 0.0022 | ns | p = 0.0022 | p = 0.0022 | p = 0.0022 |
| 52DIV | ns | p = 0.0022 | ns | p = 0.0022 | p = 0.0022 | p = 0.0022 |
| 56DIV | ns | p = 0.0022 | ns | p = 0.0043 | p = 0.0022 | p = 0.0022 |
| 59DIV | ns | p = 0.0022 | p = 0.0022 | p = 0.0022 | p = 0.0022 | p = 0.0022 |
| 63DIV | ns | p = 0.0022 | p = 0.0043 | p = 0.0022 | p = 0.0022 | p = 0.0022 |

B

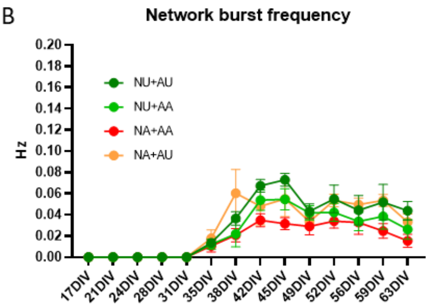


| | NU+AU vs NU+AA | NU+AA vs NA+AA | NA+AA vs NA+AU | NU+AU vs NA+AU | NU+AU vs NA+AA | NU+AA vs NA+AU |
|---|---|---|---|---|---|---|
| 17DIV | ns | ns | ns | ns | ns | ns |
| 21DIV | ns | ns | ns | ns | ns | ns |
| 24DIV | ns | ns | ns | ns | ns | ns |
| 28DIV | ns | ns | ns | ns | ns | ns |
| 31DIV | ns | ns | ns | ns | ns | ns |
| 35DIV | ns | ns | ns | ns | ns | ns |
| 38DIV | p = 0.0368 | ns | p = 0.0238 | ns | p = 0.0043 | p = 0.0476 |
| 42DIV | p = 0.0238 | p = 0.0043 | ns | ns | p = 0.0022 | ns |
| 45DIV | p = 0.0043 | p = 0.0022 | p = 0.0022 | ns | p = 0.0022 | ns |
| 49DIV | ns | p = 0.0281 | ns | p = 0.0173 | p = 0.0260 | ns |
| 52DIV | ns | ns | p = 0.0022 | ns | p = 0.0022 | p = 0.0238 |
| 56DIV | ns | ns | p = 0.0216 | ns | p = 0.0455 | p = 0.0043 |
| 59DIV | ns | ns | p = 0.0022 | ns | p = 0.0043 | p = 0.0238 |
| 63DIV | p = 0.0065 | p = 0.0390 | p = 0.0087 | ns | p = 0.0022 | ns |

C

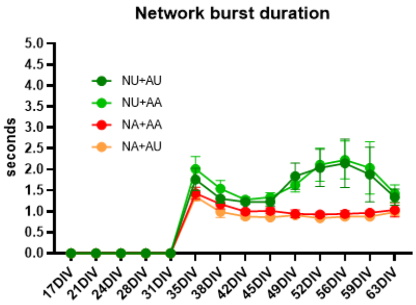


| | NU+AU vs NU+AA | NU+AA vs NA+AA | NA+AA vs NA+AU | NU+AU vs NA+AU | NU+AU vs NA+AA | NU+AA vs NA+AU |
|---|---|---|---|---|---|---|
| 17DIV | ns | ns | ns | ns | ns | ns |
| 21DIV | ns | ns | ns | ns | ns | ns |
| 24DIV | ns | ns | ns | ns | ns | ns |
| 28DIV | ns | ns | ns | ns | ns | ns |
| 31DIV | ns | ns | ns | ns | ns | ns |
| 35DIV | ns | p = 0.0022 | ns | p = 0.0043 | p = 0.0260 | p = 0.0022 |
| 38DIV | p = 0.0260 | p = 0.0043 | ns | p = 0.0238 | p = 0.0411 | p = 0.0238 |
| 42DIV | ns | p = 0.0022 | p = 0.0152 | p = 0.0022 | p = 0.0022 | p = 0.0022 |
| 45DIV | ns | p = 0.0022 | p = 0.0022 | p = 0.0022 | p = 0.0043 | p = 0.0022 |
| 49DIV | ns | p = 0.0022 | ns | p = 0.0022 | p = 0.0022 | p = 0.0022 |
| 52DIV | ns | p = 0.0022 | p = 0.0087 | p = 0.0022 | p = 0.0022 | p = 0.0022 |
| 56DIV | ns | p = 0.0022 | p = 0.0260 | p = 0.0022 | p = 0.0022 | p = 0.0022 |
| 59DIV | ns | p = 0.0022 | p = 0.0022 | p = 0.0022 | p = 0.0022 | p = 0.0022 |
| 63DIV | ns | p = 0.0022 | ns | p = 0.0043 | p = 0.0260 | p = 0.0022 |

**Supplemental Figure 9:** Development of MFR (A), NBF (B) and NBD (C) in co-cultures with neurons and astrocytes from UT (NU+AU), neurons from UT and astrocytes from AT (NU+AA), neurons and astrocytes from AT (NA+AA), and neurons from AT and astrocytes from UT (NA+AU). (n = 6 samples per condition, data was collected from 1 experiment. Mann-Whitney U test was used for the statistical comparisons)
